# The collective behavior of ant groups depends on group genotypic composition

**DOI:** 10.1101/2020.12.16.423107

**Authors:** Justin T. Walsh, Anna Garonski, Claire Jackan, Timothy A. Linksvayer

## Abstract

Increasingly, researchers document variation between groups in collective behavior, but the genetic architecture of collective behavior and how the genotypic composition of groups affects collective behavior remains unclear. Social insects are ideal for studying the effects of genetic variation on collective behavior because their societies are defined by social interactions. To explore how the genetic composition of groups affects collective behavior, we constructed groups of pharaoh ants (*Monomorium pharaonis*) from 33 genetically distinct colonies of known pedigree. The groups consisted of either all workers from the same single colony or workers from two genetically different colonies, and we assayed the exploration and aggression of the groups. We found that collective behavior depended on the specific genotypic combination of group members, i.e. we found evidence for genotype-by-genotype epistasis for both collective behaviors. Furthermore, the observed collective behavior of groups differed from the additive genetic expectations of groups, further demonstrating the importance of genotype-by-genotype effects. Finally, the collective aggression of the groups was negatively correlated with the pairwise relatedness estimates between workers within the group. Overall, this study highlights that specific combinations of genotypes influence group-level phenotypes and the difficulty of predicting group-level phenotypes using only additive models.

## Introduction

Recent research on behavior has focused on documenting behavioral variation between individuals that is consistent across time/contexts (i.e. “animal personality”; [1]). To date, consistent behavioral variation has been documented across a wide range of species and is considered ubiquitous. Similar to individuals, groups can also exhibit consistent variation in collective behavior (i.e. collective or group personality; [2–5]. For example, harvester ant (*Pogonomyrmex barbatus*) colonies differ consistently across years in their regulation of foraging rate [6–9]. Although there has been a large focus on understanding mechanistically how local behavioral interactions between individuals produce emergent group-level behavior [10], the genetic architecture of collective behavior remains largely unknown [11], including the degree to which it is heritable and how genetic variation within and between groups contributes to population-level variation for collective behavior.

Additionally, it is unclear exactly how variation in individual-level traits leads to variation in group-level traits [12–14]. For example, it is not clear whether each group member contributes equally in determining group-level behavior or if instead some individuals have a disproportionately large effect (i.e. keystone individuals). Furthermore, it is unclear how the genotypic make-up of groups affects group-level traits. For example, we do not know if individuals of a particular genotype have a consistent impact on the collective behavior of the group regardless of the genotype of other group members, or if the effect of the genotype of group members depends on the specific genotype of other group members (i.e. whether there are genotype-by-genotype interactions for collective behavior). Understanding how the genotypes of group members map to group-level phenotypes is evolutionarily important because there is increasing evidence that group-level behavioral variation influences the fitness of groups and the genetic architecture of group-level traits can affect the trait evolutionary dynamics [11, 15–19].

Social insects are ideal for elucidating the complex relationship between individual and group-level variation because they exhibit behavioral variation at multiple scales of organization (i.e. between workers, castes, colonies, species; [3]). Variation between individuals within a colony results in a division of labor, where queens reproduce while workers perform all other tasks, including foraging for food and caring for larvae [20–22]. Workers can further specialize on a wide range of tasks, resulting in division of labor within the worker caste [21–25]. Social insects exhibit consistent colony-level variation for a wide range of collective behaviors, including aggression, foraging, and exploration (reviewed by [3, 4, 26].

There is increasing evidence that colony-level behavioral variation is influenced by genotype as numerous studies have estimated the heritability of social insect collective behavior [11, 17, 27–30] and candidate gene studies have linked allelic variation to variation in collective behavior [31–34]. The amount of genetic variation within social insect colonies depends on whether the colony includes one (monogyny) or multiple queens (polygyny), whether queen(s) mates with one (monandry) or multiple males (polyandry), and the amount of inbreeding [35–39]. Both polyandry and polygyny are widespread and are thought to be favored because genetic variation within a colony may increase disease resistance [40–45] or the efficiency of division of the labor [38, 46].

A small number of studies have attempted to better understand how behavioral variation in workers influences colony-level behavioral variation by creating mixed groups that included workers with different behavioral tendencies. In these cases, one behavioral type exhibited “behavioral dominance” over the other and caused individuals within the group to behave more similarly to the dominant type. For example, in mixed colonies of docile European honey bees and aggressive Africanized honey bees, honey bees increased their aggression with the number of Africanized bees in the colony [47]. Similarly, fire ant workers will accept either one or multiple queens based on their genotype in the *Gp-9* non-recombining region, and workers from monogyne (one queen) colonies will accept multiple queens if just 5-10% of the colony consists of workers from polygyne colonies [48, 49].

In order to better understand the factors shaping collective behavior, we have to understand how the genotypic composition of the group affects it. In general, few studies have utilized an experimental design creating mixed groups and assaying continuous, rather than discrete, behavioral variation, including in social insects (but see [27, 32, 50–55] for studies on non-behavioral traits), which would allow us to understand how the genotypic composition of the group affects social insect collective behavior [56–61]. When individuals interact within a group, they can affect each other’s traits and group-level traits in an additive (i.e. G + G) or non-additive manner (i.e. G x G) [62, 63]. Non-additive effects between interacting individuals have been called G x G epistasis because epistatic interactions can exist between loci in the genomes of two interacting individuals [62–65]. Such additive and non-additive interaction effects are predicted to play an especially large role in the evolution of behavior because behavior, more so than other phenotypes, is flexible, depending on biotic and abiotic environmental conditions [66]. Furthermore, these effects are predicted to play a larger role in the evolution of behavior in social insects because of their highly complex societies that rely on social interactions [27, 67, 68].

In this study we used pharaoh ant (*Monomorium pharaonis*) colonies from a pedigreed laboratory population. Previous work on colonies from this same population found that colonies consistently varied in collective behavior and that this variation was heritable and associated with colony productivity [11]. To explore the effect of the genotypic composition of group members on the resulting collective behavior of the group, we set up groups of workers that contained either workers all from the same colony or from two different colonies, and we assayed the aggression and exploration of these groups.

## Methods

### Experimental design

We used 33 *M. pharaonis* colonies, which we subsequently refer to as “colony genotypes”, from our heterogeneous stock mapping population. This pedigreed population was started by intercrossing eight lineages for nine generations (see [11, 69, 70] for details). For each colony, we used carbon dioxide to anesthetize the workers and carefully removed 450 worker pupae using a paint brush. We separated the worker pupae into three separate petri dishes (150 pupae per dish) and monitored the dishes daily for the eclosion of callow workers. We collected the callows and placed nine callows into separate petri dishes. Five days after the callows eclosed, we combined two groups of nine callows each, either both from the same colony (“same colony groups”) or from different colonies (“mixed groups”), to form a larger group of 18 workers that we subsequently assayed for collective behavior. We refer to the two groups of workers that made up the larger group of 18 workers as “genotype one” and “genotype two”. Experimental designs including just two, rather than multiple, genotypes/families within a group have been shown theoretically to be optimal for estimating the genetic effects of group members [71]. We were able to mix workers from different colonies because *M. pharaonis* workers show little to no aggression towards conspecifics from other colonies [72, 73]. We assayed a total of 33 colonies, and the assays were divided into six blocks that each ran for about two weeks from May to August of 2018. Within each block, containing three to six total colony genotypes, each colony was paired with itself (i.e. same colony group) and with each of the other colonies three times (i.e. three replicates for each combination). We fed all groups of workers with an agar-based synthetic diet [74] and provided water *ad libitum* via a glass tubed plugged with cotton. We kept all groups of workers on a 12:12 hour light:dark cycle and at 27 ± 1 °C and 50% relative humidity.

### Behavioral assays

Two or three days after combining the two groups of nine workers into one larger group, we assayed the exploratory and aggressive behavior of the larger groups following the protocols from Walsh et al. [11]. Because some of the ants died during the aggression assay, we always conducted the exploratory assay before the aggression assay. We conducted the exploratory assay inside of a filming box with white LED lights along the walls and a camera mounted to the top [11]. To prevent trail pheromones from previous assays influencing future assays, we covered the floor of the filming box with a poster board that we replaced between each assay. We placed the 18 workers inside a Petri dish and placed the Petri dish upside down in the middle of a circular arena in the center of the filming box and waited 5 minutes for the ants to settle down after being handled. Next, we removed the petri dish, allowing the ants to explore the entirety of the circular arena, and used the camera to record the ants exploring the arena for 10 minutes. Finally, we collected all 18 workers and returned them to their Petri dish. We analyzed the videos using the R package “trackR” (https://github.com/swarm-lab/trackR), which tracked the location of all the ants in each frame of the video. We calculated the percent of the arena explored by the groups of ants by determining how many pixels were visited at least once across all frames of the video divided by the total number of pixels inside the circular arena [11].

We began the aggression assay at least two hours after the completion of the exploratory assay. Because *M. pharaonis* workers only show transient to no aggression towards conspecifics [72, 73], we quantified aggression of the *M. pharaonis* workers towards a second species, *Tetramorium immigrans* [75]. We collected the *T. immigrans* colony on the campus of the University of Pennsylvania during May of 2018 and maintained and fed the colony using the same methods we used for the *M. pharaonis* groups. We moved the 18 *M. pharaonis* workers to a small Petri dish and 18 *T. immigrans* workers to a second small Petri dish and placed the Petri dishes upside down inside a larger petri dish. We waited five minutes to give the ants time to acclimate after being handled and then lifted the small Petri dishes, allowing the ants of the two species to interact with each other. Every five minutes for one hour, we recorded the number of *M. pharaonis* workers biting *T. immigrans* workers. We defined the aggression of the groups as the average number of *M. pharaonis* workers biting *T. immigrans* workers across all observations within an hour. We froze all *T. immigrans* workers used in the assay so that we did not reuse the same workers in subsequent assays. We only managed to record aggression data for five of the six experimental blocks because our *T. immigrans* colony started to run out of workers to use in the assays.

### Statistical analyses

We performed all analyses in R version 3.6.0 [76]. To estimate the effects of genotypes one and two individually and the additive and interaction effects between them, we conducted generalized linear mixed models (GLMMs) using the R package “lme4” [77]. We note that the distinction of genotype one or two is arbitrary and could be reversed. We included the interaction between the two genotypes as a fixed effect and the block number as a random effect. To estimate the effect size of each term included in each model, we used the “r.squaredGLMM” function of the R package “MuMIn” [78].

To better understand how the interaction between genotypes one and two affected group behavior, we used the R package “MCMCglmm” [79] to build an animal model [80]. Animal models estimate genetic parameters of phenotypes by asking how phenotypic covariance between all pairs of individuals within a pedigree is predicted by the expected genetic relatedness between individuals [81, 82]. In this study, the pedigree specified the relationships between all “individuals” (i.e. colony genotypes) included as both genotypes one and two. We used the animal models to estimate the best linear unbiased predictors (BLUPs) for each genotype, which correspond to effect size estimates for genotypes one and two. Next, we asked if the observed behavior of the group was correlated with the combined BLUP for each genotype combination using Spearman rank correlation tests. Finally, we asked if the relatedness between the two genotypes influenced group behavior by conducting Spearman rank correlation tests between the behavior (either exploration or aggression) and the pairwise relatedness estimates between genotypes one and two. We conducted two-tailed correlation tests for both aggression and exploration but also conducted a one-tailed test for aggression, with the prediction that aggression decreases as relatedness increases.

## Results

The mean levels of aggression (two-tailed t test; t = 0.60, p = 0.55) and exploration (t = 0.68, p-value = 0.50) were not different between genotypes one and two, as we would expect because the decision of referring to a colony as either genotype one or two was arbitrary. The interaction between genotype one and genotype two was significant for both aggression and exploration (**Table 1**), indicating that the genotypic makeup of the group influenced group collective behavior (**Figures 1 and 2**). Block was also significant for both aggression and exploration (**Table 1**). For exploration, we estimated the proportion of variance explained by genotype one to be 0.185, by genotype two to be 0.191, by the additive effect to be 0.283, and by the interaction effect to be 0.383. (**Table 2**). For aggression, we estimated the proportion of variance explained by genotype one to be 0.347, by genotype two to be 0.351, by the additive effect to be 0.426, and by the interaction effect to be 0.498 (**Table 2**). Note that the effects of genotypes one and two and the additive effect were estimated from models that did not include the interaction term.

**Table 1.** A summary of GLM results on exploration and aggression

| **Exploration** | $\chi^2$ | df | p |
| --- | --- | --- | --- |
| Block | 18.71 | 5 | **0.002** |
| Genotype 1 x Genotype 2 | 87.72 | 55 | **0.003** |
| **Aggression** |  |  |  |
| Block | 145.43 | 4 | **<0.001** |
| Genotype 1 x Genotype 2 | 329.39 | 41 | **<0.001** |

**Figure 1.**
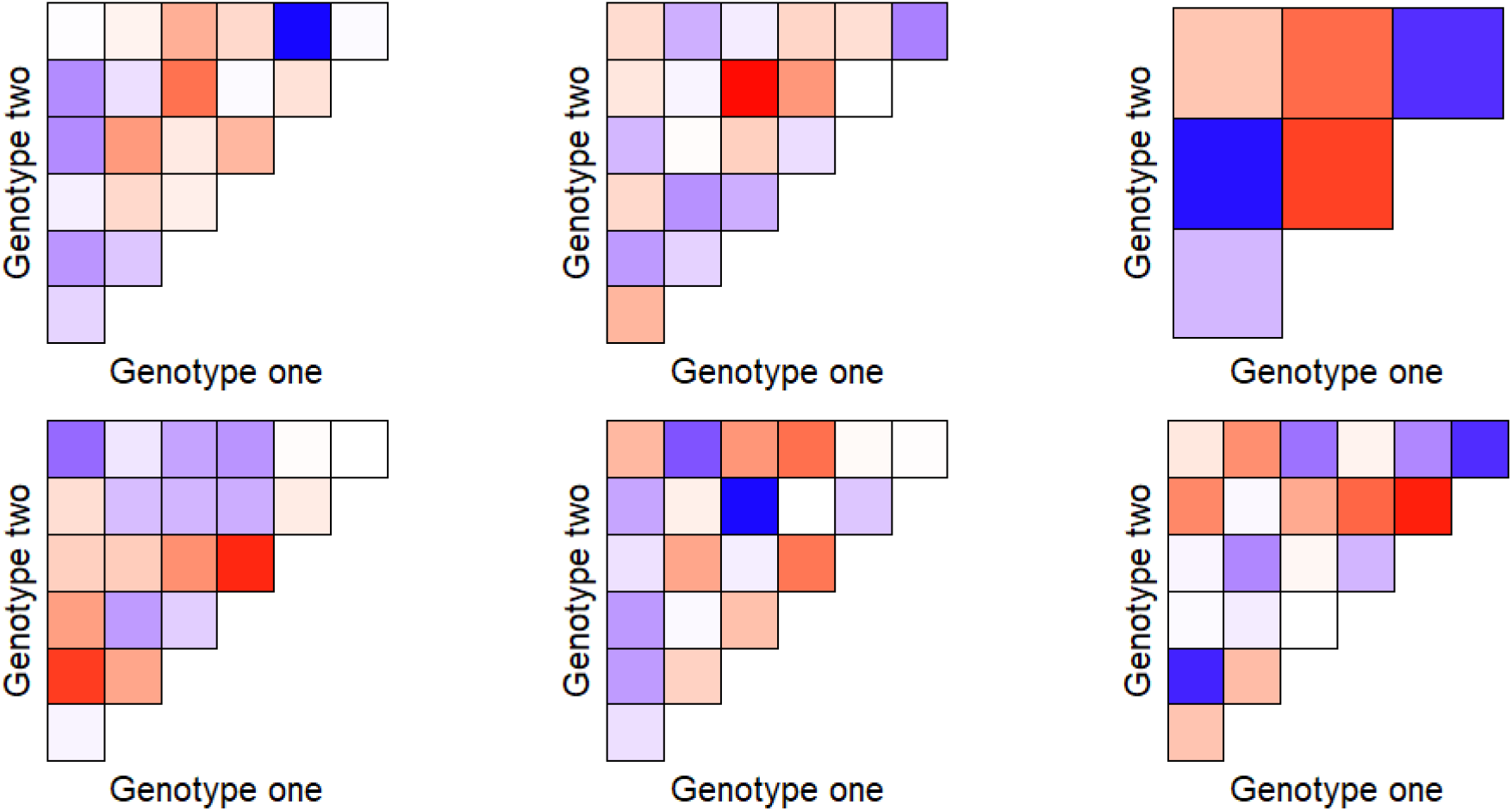
Heatmaps showing the average level of exploration between colony combinations across the six different blocks. Blue signifies high levels and red signifies low levels of aggression. Darker shades of either color signify more extreme values.

**Figure 2.**
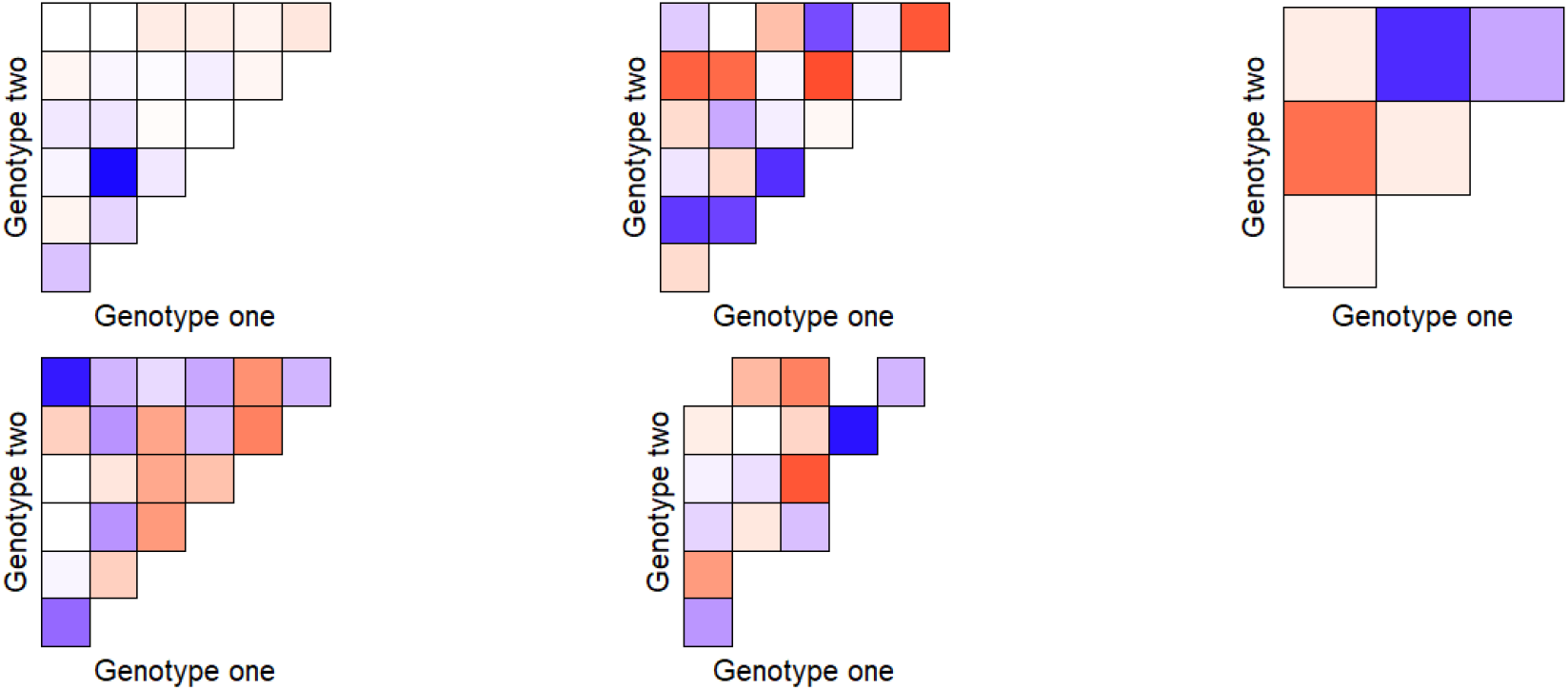
Heatmap showing the average level of aggression between colony combinations across five different blocks. Blue signifies high levels and red signifies low levels of aggression. Darker shades of either color signify more extreme values.

**Table 2.** Estimates of the contribution of DGEs, IGEs, GxG epistasis, and block effects to the total phenotypic variance for exploration and aggression

| **Exploration** | **R<sup>2</sup>** |
| --- | --- |
| Genotype one | 0.185 |
| Genotype two | 0.191 |
| G + G | 0.283 |
| G x G epistasis | 0.383 |
| Block | 0.061 |
| **Aggression** |  |
| Genotype one | 0.347 |
| Genotype two | 0.351 |
| G + G | 0.426 |
| G x G epistasis | 0.498 |
| Block | 0.236 |

We found that both the observed group exploration and aggression were significantly correlated with the expected level based on the additive combination of genotypes (as estimated as the sum of BLUPs for the two genotypes) (**Figure 3**).

**Figure 3.**
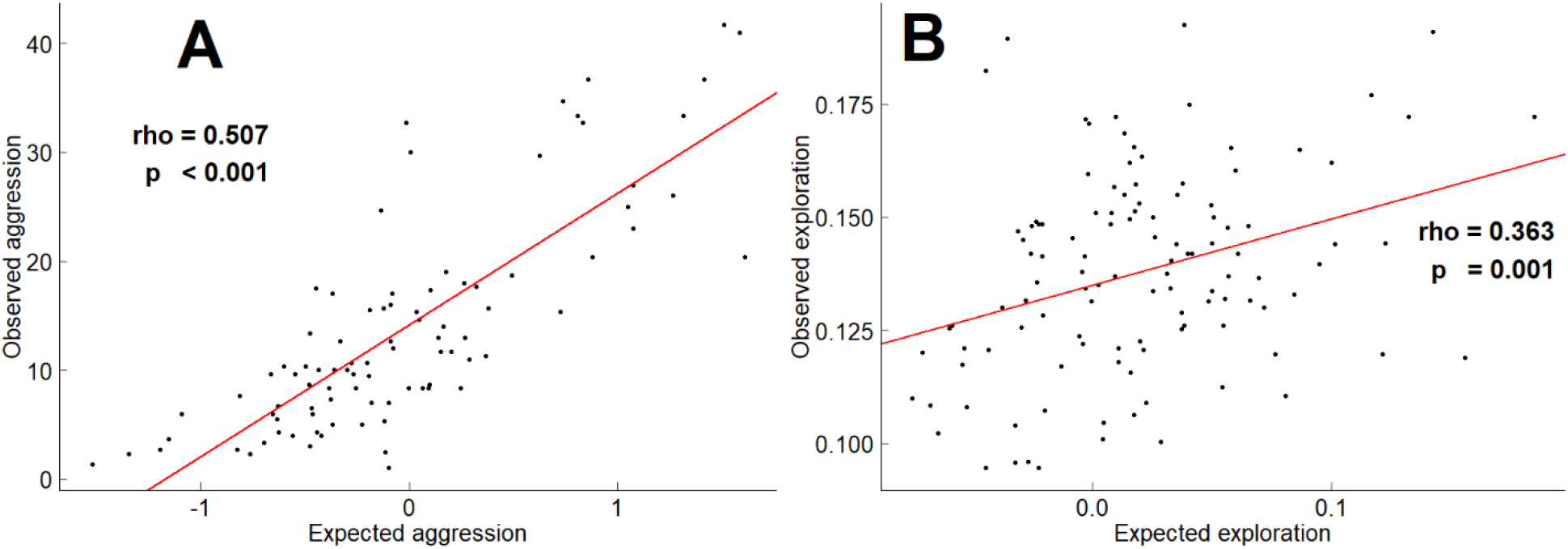
Scatterplots showing the observed group-level aggression (A) or exploration (B) compared to the expected group-level exploration, estimated as the sum of the BLUPs of genotypes one and two. The red line shows a linear model and the rho and p values are from a Spearman rank correlation test.

The pairwise pedigree relatedness between genotypes one and two was not correlated with exploration, when including the same colony groups (two-tailed Spearman rank correlation; rho = −0.035; p = 0.547) or not (rho = −0.031 ; p = 0.657), indicating that group exploration did not depend on the relatedness between group members. Furthermore, the relatedness between genotypes one and two was not correlated with aggression when including the same colony groups (two-tailed Spearman rank correlation; rho = −0.089; p = 0.151) or not (rho = −0.132 ; p = 0.072). However, we also performed a one-tailed Spearman rank correlation test for aggression because aggression might be expected to increase with within-group genetic variation (i.e. aggression to decrease as relatedness increases) due to differences in individual thresholds. We found that aggression was negatively correlated with within-group genetic relatedness when excluding the same colony groups (one-tailed Spearman rank correlation; rho = −0.133, p = 0.036).

## Discussion

The genetic architecture of group-level traits, including collective behavior, is largely unknown. We do not fully understand how the genotypic make-up of groups affects group-level traits, including to what extent additive and non-additive interactions between the genotypes of group members are important. In this study, we used a phenotypically and genetically variable laboratory population of *M. pharaonis* to study the importance of intra-group genetic composition in the production of emergent group-level behavior. In groups of ants composed of workers from two different colony genotypes, the effect of the genotype of group members was conditional on the other group members (i.e., the interaction term was significant), suggesting G x G epistasis is important. Overall, our results highlight the importance of specific genotypic combinations of group members on collective phenotypes in general.

G x G epistasis explained a larger proportion of variance than either genotype alone or the additive value between them, suggesting that the specific combinations of genotypes within a group has a large effect on variation in collective behavior (**Figures 1 and 2**). We further explored the interaction effect by estimating the effect sizes for genotypes one and two (i.e. BLUPs) while incorporating relatedness estimates based on the pedigree (**Figure 3**). The sum of the two BLUPs represents the expected phenotype if the genetic effects of interacting group members on collective behavior was additive. We specifically were interested in whether there were any obvious patterns in the relationship between observed and expected behavioral outputs. For example, we could have observed behavioral compensation, a situation where different genotypic combinations give similar phenotypic outcomes meaning that as the predicted phenotypic level increased, the observed phenotype would level off or remain flat. Although there is not an obvious pattern of deviation from the expected values (and both behaviors were significantly correlated with the combined BLUP values), there is a large amount of deviation from the expected behavioral values, further emphasizing the point that the specific combination of genotypes one and two played a large role in determining the behavior of the group.

Group-level phenotypes depend on potentially complex genetic interaction effects between group members. Therefore, simple additive models, models that either consider only additive effects or ignore the genotypes of social partners entirely, are not adequate [56, 61, 62]. When considering only a focal individual’s genotype, models assume that the covariance between genotype and phenotype is equivalent to the additive genetic variance with an individual but this is not the case when either additive effects or G x G epistasis between focal individuals and social partners are present [56, 61, 83, 84]. Furthermore, both additive effects and G x G epistasis between social partners are genetic and, therefore, represent heritable components of the environment, meaning that the environment itself can respond to selection and evolve over time [56, 61, 62]. Evolutionary models considering only genetic effects in focal individuals fail to incorporate the changes to the mean phenotype provided by the genetic components of the environment. Our results build on previous work demonstrating that when attempting to understand the genotype-phenotype relationship or the evolutionary response to selection researchers strongly consider both additive and interaction effects between social partners (reviewed by [66, 85]. Our results highlight that group-level phenotypes are difficult to predict from additive expectations.

Although social interactions, and therefore the potential for additive effects and G x G epistasis, occur between conspecifics of almost every species, these effects are predicted to be especially important for social insects [68]. Social insect colonies are characterized by a division of labor between individuals within the colony, which requires frequent communication between individuals, and by cooperative brood care by workers [21, 22, 27, 67, 68]. The occurrence of G x G epistasis likely depends on the amount of genetic variation within the colony. The amount of genetic diversity within a social insect colony depends on a number of factors including whether the colony is monogynous or polygynous, the level of polyandry, and the amount of inbreeding [35–39]. We would expect the effect of G x G epistasis on colony-level phenotypes to be larger in colonies with higher levels of within-colony genotypic variation, due to more queens, higher levels of polyandry, and low levels of inbreeding. Additionally, in unicolonial species, including *M. pharaonis*, individual workers can freely move between neighboring colonies, leading to more genetic diversity [86] and a greater potential for G x G epistasis between genotypes. Finally, genetic diversity within social insect colonies allows for “social heterosis,” the maintenance of genetic diversity through a mutualistic benefit of the inter-individual expression of multiple alleles at a single locus [87].

We found that group-level aggression, but not exploration, was negatively correlated with the pairwise relatedness between genotypes one and two within a group. While we might expect aggression towards conspecifics to increase as relatedness decreases (although unicolonial species like *M. pharaonis* show little to no aggression towards conspecifics; [72, 73]), our assay quantified aggression towards a second species (*T. immigrans*). The response-threshold model postulates that workers within a social insect colony differ intrinsically, possibly due to genotypic differences [88], in the stimulus level at which they begin to behaviorally respond [22, 89, 90]. Increased genetic diversity within a social insect group would increase the likelihood that at least some individuals would respond to a stimulus at a given level. Therefore, *M. pharaonis* group-level aggression may have increased as genetic diversity increased (as relatedness within the group decreased) because there was a greater chance that some group members would respond aggressively to the *T. immigrans* workers and recruit others, through the use of alarm pheromones, to also respond aggressively.

Overall, this study highlights the importance of the specific combinations of genotypes in shaping collective behavior. We detected an effect of G x G epistasis when only studying small groups of 18 workers, which are much less complex than typical *M. pharaonis* colonies that can include thousands of workers in addition to multiple queens and brood at different developmental stages, suggesting that these effects are widespread, as predicted. Furthermore, our small groups included only two distinct genotypes, while real colonies may have a wider range of genotypes. Additionally, our study was conducted in the laboratory, under carefully controlled environmental conditions. In a natural setting, genotype-by-environment interaction effects are likely very common, further complicating how genetic composition affects group-level traits. Future studies should tease apart how the genotypic composition of group members influences social interactions through different types of social communication (e.g. pheromones, physical interactions, trophallaxis) and how social interactions influence phenotypic variation across all colony members (e.g. queens, workers, brood). Finally, future studies should also attempt to identify specific genes that play a role in the social regulation of phenotypes. One such approach could be to examine the composition of “social fluids,” fluids passed between individuals (e.g. trophallaxis between workers or between workers and larvae), and the gene expression of the tissues that produce the fluids, as social fluids have been shown to be an important mechanism by which workers regulate the development of brood [91].

## Acknowledgements

Rohini Singh and Michael Warner provided feedback on the experimental design. Joel McGlothlin provided feedback on the statistical analysis.

## References

1. Sih A., Bell A., Johnson J.C. 2004 Behavioral syndromes: an ecological and evolutionary overview. Trends in Ecology & Evolution 19(7), 372–378. (doi:https://doi.org/10.1016/j.tree.2004.04.009).

2. Gordon D.M. 1991 Behavioral flexibility and the foraging ecology of seed-eating ants. The American Naturalist 138(2), 379–411. (doi:10.1086/285223).

3. Jandt J.M., Bengston S., Pinter-Wollman N., Pruitt J.N., Raine N.E., Dornhaus A., Sih A. 2014 Behavioural syndromes and social insects: personality at multiple levels. Biol Rev 89(1), 48–67. (doi:10.1111/brv.12042).

4. Bengston S., Jandt J. 2014 The development of collective personality: the ontogenetic drivers of behavioral variation across groups. Frontiers in Ecology and Evolution 2(81). (doi:10.3389/fevo.2014.00081).

5. Wright C.M., Lichtenstein J.L.L., Doering G.N., Pretorius J., Meunier J., Pruitt J.N. 2019 Collective personalities: present knowledge and new frontiers. Behavioral Ecology and Sociobiology 73(3), 31. (doi:10.1007/s00265-019-2639-2).

6. Gordon D.M. 2002 The regulation of foraging activity in red harvester ant colonies. The American Naturalist 159(5), 509–518. (doi:10.1086/339461).

7. Greene M.J., Gordon D.M. 2007 Interaction rate informs harvester ant task decisions. Behavioral Ecology 18(2), 451–455. (doi:10.1093/beheco/arl105).

8. Gordon D.M., Holmes S., Nacu S. 2007 The short-term regulation of foraging in harvester ants. Behavioral Ecology 19(1), 217–222. (doi:10.1093/beheco/arm125).

9. Gordon D.M., Guetz A., Greene M.J., Holmes S. 2011 Colony variation in the collective regulation of foraging by harvester ants. Behavioral Ecology 22(2), 429–435. (doi:10.1093/beheco/arq218).

10. Sumpter D.J.T. 2010 Collective animal behavior. Princeton, NJ, US, Princeton University Press.

11. Walsh J.T., Garnier S., Linksvayer T.A. 2020 Ant Collective Behavior Is Heritable and Shaped by Selection. The American Naturalist 196(5), 541–554. (doi:10.1086/710709).

12. Pinter-Wollman N. 2012 Personality in social insects: How does worker personality determine colony personality? Current Zoology 58(4), 580–588. (doi:10.1093/czoolo/58.4.580).

13. LeBoeuf A.C., Grozinger C.M. 2014 Me and we: the interplay between individual and group behavioral variation in social collectives. Current Opinion in Insect Science 5, 16–24. (doi:https://doi.org/10.1016/j.cois.2014.09.010).

14. Ulrich Y., Kawakatsu M., Tokita C.K., Saragosti J., Chandra V., Tarnita C.E., Kronauer D.J.C. 2020 Emergent behavioral organization in heterogeneous groups of a social insect. bioRxiv, 2020.2003.2005.963207. (doi:10.1101/2020.03.05.963207).

15. Wray M.K., Mattila H.R., Seeley T.D. 2011 Collective personalities in honeybee colonies are linked to colony fitness. Animal Behaviour 81(3), 559–568. (doi:https://doi.org/10.1016/j.anbehav.2010.11.027).

16. Modlmeier A.P., Liebmann J.E., Foitzik S. 2012 Diverse societies are more productive: a lesson from ants. Proceedings of the Royal Society B: Biological Sciences 279(1736), 2142–2150. (doi:10.1098/rspb.2011.2376).

17. Gordon D.M. 2013 The rewards of restraint in the collective regulation of foraging by harvester ant colonies. Nature 498, 91. (doi:10.1038/nature12137 https://www.nature.com/articles/nature12137#supplementary-information).

18. Blight O., Albet Díaz-Mariblanca G., Cerdá X., Boulay R. 2016 A proactive–reactive syndrome affects group success in an ant species. Behavioral Ecology 27(1), 118–125. (doi:10.1093/beheco/arv127).

19. Blight O., Villalta I., Cerdá X., Boulay R. 2016 Personality traits are associated with colony productivity in the gypsy ant *Aphaenogaster senilis*. Behav Ecol Sociobiol 70, 2203–2209 (2016). https://doi.org/10.1007/s00265-016-2224-x

20. Wilson EO. 1971 The insect societies. Cambridge, MA: Belknap Press of Harvard University Press.

21. Oster G.F., Wilson E.O. 1978 Caste and ecology in the social insects. Monographs in population biology 12, 1–352.

22. Beshers S.N., Fewell J.H. 2001 Models of division of labor in social insects. Annu Rev Entomol 46, 413–440. (doi:10.1146/annurev.ento.46.1.413).

23. Robinson G.E. 1992 Regulation of Division of Labor in Insect Societies. Annual Review of Entomology 37(1), 637–665. (doi:10.1146/annurev.en.37.010192.003225).

24. Mikheyev A.S., Linksvayer T.A. 2015 Genes associated with ant social behavior show distinct transcriptional and evolutionary patterns. eLife 4, e04775. (doi:10.7554/eLife.04775).

25. Walsh J.T., Warner M.R., Kase A., Cushing B.J., Linksvayer T.A. 2018 Ant nurse workers exhibit behavioural and transcriptomic signatures of specialization on larval stage. Animal Behaviour 141, 161–169. (doi:https://doi.org/10.1016/j.anbehav.2018.05.015).

26. Jeanson R., Weidenmüller A. 2014 Interindividual variability in social insects – proximate causes and ultimate consequences. Biol Rev 89(3), 671–687. (doi:https://doi.org/10.1111/brv.12074).

27. Linksvayer T.A. 2006 Direct, maternal, and sibsocial genetic effects on individual and colony traits in an ant. Evolution 60(12), 2552–2561. (doi:10.1111/j.0014-3820.2006.tb01889.x).

28. Hunt G.J., Amdam G.V., Schlipalius D., Emore C., Sardesai N., Williams C.E., Rueppell O., Guzmán-Novoa E., Arechavaleta-Velasco M., Chandra S., et al. 2007 Behavioral genomics of honeybee foraging and nest defense. Naturwissenschaften 94(4), 247–267. (doi:10.1007/s00114-006-0183-1).

29. Greenwood A.K., Ardekani R., McCann S.R., Dubin M.E., Sullivan A., Bensussen S., Tavaré S., Peichel C.L. 2015 Genetic mapping of natural variation in schooling tendency in the threespine stickleback. G3: Genes|Genomes|Genetics 5(5), 761–769. (doi:10.1534/g3.114.016519).

30. Friedman D.A., Gordon D.M. 2016 Ant genetics: reproductive physiology, worker morphology, and behavior. Annual Review of Neuroscience 39(1), 41–56. (doi:10.1146/annurev-neuro-070815-013927).

31. Krieger M.J. 2005 To b or not to b: a pheromone-binding protein regulates colony social organization in fire ants. BioEssays : news and reviews in molecular, cellular and developmental biology 27(1), 91–99. (doi:10.1002/bies.20129).

32. Wang J., Ross K.G., Keller L. 2008 Genome-wide expression patterns and the genetic architecture of a fundamental social trait. PLOS Genetics 4(7), e1000127. (doi:10.1371/journal.pgen.1000127).

33. Wang J., Wurm Y., Nipitwattanaphon M., Riba-Grognuz O., Huang Y.-C., Shoemaker D., Keller L. 2013 A Y-like social chromosome causes alternative colony organization in fire ants. Nature 493, 664. (doi:10.1038/nature11832 https://www.nature.com/articles/nature11832#supplementary-information).

34. Tang W., Zhang G., Serluca F., Li J., Xiong X., Coble M., Tsai T., Li Z., Molind G., Zhu P., et al. 2018 Genetic architecture of collective behaviors in zebrafish. bioRxiv, 350314. (doi:10.1101/350314).

35. Keller L. 1993 Queen number and sociality in insects, Oxford University Press Oxford.

36. Bourke A.F., Franks N.R. 1995 Social evolution in ants, Princeton University Press.

37. Boomsma J.J., Ratnieks F.L. 1996 Paternity in eusocial Hymenoptera. Philosophical Transactions of the Royal Society of London Series B: Biological Sciences 351(1342), 947–975.

38. Oldroyd B.P., Fewell J.H. 2007 Genetic diversity promotes homeostasis in insect colonies. Trends in ecology & evolution 22(8), 408–413.

39. Haag-Liautard C., Vitikainen E., Keller L., Sundström L. 2009 Fitness and the level of homozygosity in a social insect. J Evol Biol 22(1), 134–142. (doi:10.1111/j.1420-9101.2008.01635.x).

40. Baer B., Schmid-Hempel P. 1999 Experimental variation in polyandry affects parasite loads and fitness in a bumble-bee. Nature 397(6715), 151–154. (doi:10.1038/16451).

41. Schmid-Hempel P., Crozier R.H. 1999 Ployandry versus polygyny versus parasites. Philos Trans R Soc Lond B Biol Sci 354(1382), 507–515. (doi:10.1098/rstb.1999.0401).

42. Tarpy D.R. 2003 Genetic diversity within honeybee colonies prevents severe infections and promotes colony growth. Proceedings of the Royal Society B: Biological sciences 270(1510), 99–103. (doi:10.1098/rspb.2002.2199).

43. Hughes W.O., Boomsma J.J. 2004 Genetic diversity and disease resistance in leaf◻cutting ant societies. Evolution 58(6), 1251–1260.

44. Tarpy D.R., Seeley T.D. 2006 Lower disease infections in honeybee (*Apis mellifera*) colonies headed by polyandrous vs monandrous queens. Naturwissenschaften 93(4), 195–199. (doi:10.1007/s00114-006-0091-4).

45. Reber A., Castella G., Christe P., Chapuisat M. 2008 Experimentally increased group diversity improves disease resistance in an ant species. Ecology letters 11(7), 682–689. (doi:10.1111/j.1461-0248.2008.01177.x).

46. Crozier R.H., Page R.E. 1985 On being the right size: male contributions and multiple mating in social Hymenoptera. Behavioral Ecology and Sociobiology 18(2), 105–115. (doi:10.1007/BF00299039).

47. Guzmán-Novoa E., Page R.E., Jr. 1994 Genetic dominance and worker interactions affect honeybee colony defense. Behavioral Ecology 5(1), 91–97. (doi:10.1093/beheco/5.1.91).

48. Ross K., Keller L. 2002 Experimental conversion of colony social organization by manipulation of worker genotype composition in fire ants (*Solenopsis invicta*). Behavioral Ecology and Sociobiology 51(3), 287–295.

49. Gotzek D., Ross K.G. 2008 Experimental conversion of colony social organization in fire ants (*Solenopsis invicta*): worker genotype manipulation in the absence of queen effects. Journal of Insect Behavior 21(5), 337–350. (doi:10.1007/s10905-008-9130-7).

50. Linksvayer T.A. 2007 Ant species differences determined by epistasis between brood and worker genomes. PLOS ONE 2(10), e994. (doi:10.1371/journal.pone.0000994).

51. Linksvayer Timothy A., Fondrk Michael K., Page Jr Robert E. 2009 Honeybee social regulatory networks are shaped by colony_Jlevel selection. The American Naturalist 173(3), E99–E107. (doi:10.1086/596527).

52. Linksvayer T.A., Kaftanoglu O., Akyol E., Blatch S., Amdam G.V., Page Jr R.E. 2011 Larval and nurse worker control of developmental plasticity and the evolution of honey bee queen–worker dimorphism. Journal of Evolutionary Biology 24(9), 1939–1948. (doi:10.1111/j.1420-9101.2011.02331.x).

53. Teseo S., Châline N., Jaisson P., Kronauer D.J. 2014 Epistasis between adults and larvae underlies caste fate and fitness in a clonal ant. Nature communications 5(1), 1–8.

54. van Zweden J.S., Brask J.B., Christensen J.H., Boomsma J.J., Linksvayer T.A., d’Ettorre P. 2010 Blending of heritable recognition cues among ant nestmates creates distinct colony gestalt odours but prevents within-colony nepotism. Journal of Evolutionary Biology 23(7), 1498–1508. (doi:10.1111/j.1420-9101.2010.02020.x).

55. Vojvodic S., Johnson B.R., Harpur B.A., Kent C.F., Zayed A., Anderson K.E., Linksvayer T.A. 2015 The transcriptomic and evolutionary signature of social interactions regulating honey bee caste development. Ecology and evolution 5(21), 4795–4807. (doi:10.1002/ece3.1720).

56. Moore A.J., Brodie Iii E.D., Wolf J.B. 1997 Interacting phenotypes and the evolutionary process: I. Direct and indirect genetic effects of social interations. Evolution 51(5), 1352–1362. (doi:10.1111/j.1558-5646.1997.tb01458.x).

57. Bijma P., Muir W.M., Van Arendonk J.A.M. 2007 Multilevel selection 1: Quantitative genetics of inheritance and response to selection. Genetics 175(1), 277. (doi:10.1534/genetics.106.062711).

58. Bijma P., Muir W.M., Ellen E.D., Wolf J.B., Van Arendonk J.A.M. 2007 Multilevel selection 2: Estimating the genetic parameters determining inheritance and response to selection. Genetics 175(1), 289. (doi:10.1534/genetics.106.062729).

59. Bijma P., Wade M.J. 2008 The joint effects of kin, multilevel selection and indirect genetic effects on response to genetic selection. Journal of evolutionary biology 21(5), 1175–1188.

60. Bijma P. 2011 A general definition of the heritable variation that determines the potential of a population to respond to selection. Genetics 189(4), 1347–1359. (doi:10.1534/genetics.111.130617).

61. McGlothlin J.W., Moore A.J., Wolf J.B., Brodie III E.D. 2010 Interacting phenotypes and the evolutionary process. III. Social evolution. Evolution 64(9), 2558–2574. (doi:doi:10.1111/j.1558-5646.2010.01012.x).

62. Wolf J.B., Brodie Iii E.D., Cheverud J.M., Moore A.J., Wade M.J. 1998 Evolutionary consequences of indirect genetic effects. Trends Ecol Evol 13(2), 64–69.

63. Wade M.J. 2000 Opposing levels of selection can cause neutrality; mating patterns and maternal-fetal interactions. Evolution 54(1), 290–292. (doi:10.1111/j.0014-3820.2000.tb00029.x).

64. Culumber Z.W., Kraft B., Lemakos V., Hoffner E., Travis J., Hughes K.A. 2018 GxG epistasis in growth and condition and the maintenance of genetic polymorphism in *Gambusia holbrooki*. Evolution 72(5), 1146–1154. (doi:https://doi.org/10.1111/evo.13474).

65. Jaffe A., Burns M.P., Saltz J.B. 2020 Genotype-by-genotype epistasis for exploratory behaviour in *D. simulans*. Proceedings of the Royal Society B: Biological Sciences 287(1928), 20200057. (doi:doi:10.1098/rspb.2020.0057).

66. Bailey N.W., Marie-Orleach L., Moore A.J. 2017 Indirect genetic effects in behavioral ecology: does behavior play a special role in evolution? Behavioral Ecology 29(1), 1–11. (doi:10.1093/beheco/arx127).

67. Linksvayer T.A., Wade M.J. 2005 The Evolutionary Origin and Elaboration of Sociality in the Aculeate Hymenoptera: Maternal Effects, Sib◻Social Effects, and Heterochrony. The Quarterly Review of Biology 80(3), 317–336. (doi:10.1086/432266).

68. Linksvayer T.A. 2015 The Molecular and Evolutionary Genetic Implications of Being Truly Social for the Social Insects. In Genomics, Physiology and Behaviour of Social Insects (eds. Zayed A., Kent C.F.), pp. 271–292. London, Academic Press Ltd-Elsevier Science Ltd.

69. Walsh J., Pontieri L., d’Ettorre P., Linksvayer T.A. 2020 Ant cuticular hydrocarbons are heritable and associated with variation in colony productivity. Proceedings of the Royal Society B: Biological Sciences 287(1928), 20201029. (doi:doi:10.1098/rspb.2020.1029).

70. Pontieri L., Schmidt A.M., Singh R., Pedersen J.S., Linksvayer T.A. 2017 Artificial selection on ant female caste ratio uncovers a link between female-biased sex ratios and infection by *Wolbachia* endosymbionts. Journal of Evolutionary Biology 30(2), 225–234. (doi:10.1111/jeb.13012).

71. Bijma P. 2010 Estimating indirect genetic effects: precision of estimates and optimum designs. Genetics 186(3), 1013–1028. (doi:10.1534/genetics.110.120493).

72. Schmidt A.M., d’Ettorre P., Pedersen J.S. 2010 Low levels of nestmate discrimination despite high genetic differentiation in the invasive pharaoh ant. Frontiers in Zoology 7(1), 20. (doi:10.1186/1742-9994-7-20).

73. Pontieri L. 2014 Discrimination behavior in the supercolonial pharaoh ant. PhD thesis, University of Copenhagen, København, Denkmark

74. Dussutour A., Simpson S.J. 2008 Description of a simple synthetic diet for studying nutritional responses in ants. Insectes Sociaux 55(3), 329–333. (doi:10.1007/s00040-008-1008-3).

75. Wagner H.C., Arthofer W., Seifert B., Muster C., Steiner F.M., Schlick-Steiner B.C. 2017 Light at the end of the tunnel: Integrative taxonomy delimits cryptic species in the *Tetramorium caespitum* complex (Hymenoptera: Formicidae). Myrmecological News 25, 95–129. (doi:10.25849/myrmecol.news_025:095).

76. R Core Team. 2013 R: a language and environment for statistical computing. Vienna, Austria: R Foundation for Statistical Computing.

77. Bates D, Mächler M, Bolker B, Walker S (2015). “Fitting Linear Mixed-Effects Models Using lme4.” Journal of Statistical Software, 67(1), 1–48. doi: 10.18637/jss.v067.i01.

78. Barton, K. (2009) Mu-MIn: Multi-model inference. R Package Version 0.12.2/r18. http://R-Forge.R-project.org/projects/mumin/

79. Hadfield J.D. 2010 MCMC methods for multi-response generalized linear mixed models: The MCMCglmm R package. 2010 33(2), 22. (doi:10.18637/jss.v033.i02).

80. Wilson A.J., Reale D., Clements M.N., Morrissey M.M., Postma E., Walling C.A., Kruuk L.E., Nussey D.H. 2010 An ecologist’s guide to the animal model. The Journal of animal ecology 79(1), 13–26. (doi:10.1111/j.1365-2656.2009.01639.x).

81. Kruuk L.E.B. 2004 Estimating genetic parameters in natural populations using the “animal model”. Philosophical transactions of the Royal Society of London Series B, Biological sciences 359(1446), 873–890. (doi:10.1098/rstb.2003.1437).

82. de Villemereuil, P. 2012. Tutorial estimation of a biological trait heritability using the animal model: how to use the MCMCglmm R package. https://devillemereuil.legtux.org/wp-content/uploads/2012/12/tuto_en.pdf

83. Kirkpatrick M., Lande R. 1989 The evolution of maternal characters. Evolution 43(3), 485–503. (doi:10.2307/2409054).

84. Lande R., Kirkpatrick M. 1990 Selection response in traits with maternal inheritance. Genetical research 55(3), 189–197. (doi:10.1017/s0016672300025520).

85. Bijma P. 2014 The quantitative genetics of indirect genetic effects: a selective review of modelling issues. Heredity (Edinb) 112(1), 61–69. (doi:10.1038/hdy.2013.15).

86. Giraud T., Pedersen J.S., Keller L. 2002 Evolution of supercolonies: The Argentine ants of southern Europe. Proceedings of the National Academy of Sciences 99(9), 6075–6079. (doi:10.1073/pnas.092694199).

87. Nonacs P., Kapheim K.M. 2007 Social heterosis and the maintenance of genetic diversity. Journal of Evolutionary Biology 20(6), 2253–2265. (doi:10.1111/j.1420-9101.2007.01418.x).

88. Page R.E., Robinson G.E. 1991 The genetics of division of labour in honey bee colonies. In Advances in Insect Physiology (ed. Evans P.D.), pp. 117–169, Academic Press.

89. Wilson E.O. 1976 Behavioral discretization and the number of castes in an ant species. Behavioral Ecology and Sociobiology 1(2), 141–154. (doi:10.1007/BF00299195).

90. Robinson G.E. 1987 Regulation of honey bee age polyethism by juvenile hormone. Behavioral Ecology and Sociobiology 20(5), 329–338. (doi:10.1007/BF00300679).

91. LeBoeuf A.C., Waridel P., Brent C.S., Gonçalves A.N., Menin L., Ortiz D., Riba-Grognuz O., Koto A., Soares Z.G., Privman E., et al. 2016 Oral transfer of chemical cues, growth proteins and hormones in social insects. eLife 5, e20375. (doi:10.7554/eLife.20375).

